# Ultra-deep long-read metagenomics captures diverse taxonomic and biosynthetic potential of soil microbes

**DOI:** 10.1101/2025.05.28.656579

**Authors:** Caner Bağcı, Timo Negri, Elena Buena Atienza, Caspar Gross, Stephan Ossowski, Nadine Ziemert

**Affiliations:** Translational Genome Mining for Natural Products, Interfaculty Institute of Microbiology and Infection Medicine Tübingen (IMIT), University of Tübingen, Auf der Morgenstelle 24, 72076, Tübingen, Baden-Württemberg, Germany; Interfaculty Institute for Biomedical Informatics (IBMI), University of Tübingen, Auf der Morgenstelle 24, 72076, Tübingen, Baden-Württemberg, Germany; German Center for Infection Research (DZIF), Partner Site Tübingen, Auf der Morgenstelle 24, 72076, Tübingen, Baden-Württemberg, Germany; Institute of Medical Genetics and Applied Genomics, University of Tübingen, Calwerstrasse 7, 72076, Tübingen, Baden-Württemberg, Germany; NGS Competence Center, University of Tübingen, Calwerstrasse 7, 72076, Tübingen, Baden-Württemberg, Germany

**Keywords:** microbiome, soil, metagenomics, nanopore, ultra-deep, long-read sequencing, diversity, natural products, (meta)genome mining

## Abstract

**Background:** Soil ecosystems have long been recognised as hotspots of microbial diversity, but most estimates of their microbial and functional complexity remain speculative despite decades of study, in part because conventional sequencing campaigns lack the depth and contiguity required to recover low-abundance and repetitive genomes. Here, we revisit this question using one of the deepest metagenomic sequencing efforts to date, applying 148 billion base pairs of Nanopore long-read and 122 billion base pairs of Illumina short-read data to a single forest soil sample.

**Results:** Our hybrid assembly reconstructed 837 metagenome-assembled genomes, including 466 that meet high- and medium-quality standards, nearly all lacking close relatives among cultivated taxa. Rarefaction and k-mer analyses reveal that, even at this depth, we capture only a fraction of the extant diversity: non-parametric models project that more than ten trillion base pairs of sequencing data would be required to approach saturation. These findings offer a quantitative, technology-enabled update to long-standing diversity estimates and demonstrate that conventional metagenomic sequencing efforts likely miss the majority of microbial and biosynthetic potential in soil. We further identify more than 11 000 biosynthetic gene clusters, over 99% of which have no match in current databases, underscoring the breadth of unexplored metabolic capacity.

**Conclusions:** Taken together, our results emphasise both the power and the present limitations of metagenomics in resolving natural microbial complexity, and they provide a new baseline for evaluating future advances in microbial genome recovery, taxonomic classification, and natural product discovery.

**Data Description:** In order to quantify how much taxonomic and biosynthetic novelty present-day sequencing can recover from soil, we extracted high-molecular-weight DNA from a single Cambisol forest soil sample (Schönbuch, Germany) and generated an ultra-deep 270 Gbp dataset - 148 Gbp of Oxford Nanopore PromethION long reads (read-length N50 = 12.2 kb) plus 122 Gbp of Illumina NovaSeq reads. The hybrid metagenome assembly we conducted produced a 10.5 Gbp assembly, from which multi-tool binning and subsequent refinement recovered 837 MAGs, and antiSMASH/BiG-SCAPE annotation revealed more than 11 000 largely novel biosynthetic gene clusters, creating a resource for benchmarking assembly or binning pipelines, modelling diversity-coverage relationships and mining natural products; all raw reads are archived under ENA BioProject PRJEB89893 [1], and the polished assembly, MAG set and BGC catalogue are available via Zenodo for unrestricted reuse [2, 3].

**Key Points:**

- Ultra-deep hybrid sequencing (148 Gbp Nanopore + 122 Gbp Illumina) of a single forest soil sample yielded 837 metagenome-assembled genomes, all lacking cultured counterparts.
- Despite this unprecedented 270 Gbp depth, rarefaction and coverage modelling indicate that more than 10 Tbp of data would still be needed to reach saturation in soil.
- Only 0.7% and 16.7% of all assembled contigs can be assigned to a species and genus, respectively, with at least one cultured representative, highlighting an unprecedented level of novelty in soil.
- The assembly uncovers 11 381 biosynthetic gene clusters forming over 10 000 mostly novel families, spotlighting an immense, untapped reservoir of microbial natural-product potential.

## Introduction

Soil is one of the most biologically diverse ecosystems on Earth, hosting a massive and largely uncharacterised diversity of microbial life. These microbial communities play critical roles in biogeochemical cycling, soil formation, nutrient turnover, plant health, and climate regulation [4]. Despite their ecological and biotechno-logical importance, the full extent and functional potential of soil microbiomes remain poorly understood.

A single gram of soil can contain up to 10^9^ microbial cells and hundreds of thousands of species, spanning all domains of life [5, 6]. These communities are not only taxonomically complex but also exhibit highly uneven abundance distributions, where rare taxa – collectively termed the “rare biosphere” – may be functionally consequential despite their low abundance [7].

Although the vast diversity of soil microbes has been widely acknowledged for decades, most estimates of their richness are based on indirect approaches, including 16S rRNA gene surveys, limited shotgun datasets, or predictive modelling. As a result, foundational claims about the extent of soil microbial diversity are frequently cited but rarely re-examined with contemporary data. Much of the soil microbiome remains part of the so-called microbial dark matter: lineages with no cultured representatives and no reference genomes [8, 9, 10].

Metagenomics has enabled the cultivation-independent study of microbial communities, allowing for genome-resolved insights into uncultured organisms [11, 12, 13]. However, most metagenomic studies rely on short-read sequencing, which presents challenges in assembling highly complex, strain-rich communities like soil. These limitations hinder the recovery of low-abundance taxa, reduce the completeness of biosynthetic gene clusters, and often lead to fragmented or ambiguous assemblies [12]. Recent advances in long-read sequencing, particularly with Oxford Nanopore Technologies (ONT), have opened new opportunities for soil microbiome research. Long reads can span repeat-rich regions and operons, improving genome recovery and assembly contiguity [14, 15, 16]. Hybrid approaches combining long and short reads can further mitigate the high error rates of long reads while leveraging their structural advantages [17, 18]. Despite these improvements, it remains unclear how far even the most advanced sequencing technologies can go in capturing the full taxonomic and functional diversity of soil.

To establish an empirical reference point for assessing microbial and biosynthetic diversity in soil, we performed one of the deepest metagenomic sequencing efforts to date on a single soil sample, combining 148 billion base pairs (Gbp) of ONT long-read and 122 Gbp of Illumina short-read data. We use this dataset to empirically assess the power and current limitations of metagenomics in resolving complex microbial communities. Specifically, we ask: How much taxonomic and biosynthetic diversity can be recovered from a single soil sample using state-of-the-art sequencing? What proportion of this diversity is represented in existing databases? Can the ultra-deep sequencing approach saturation, or are we still only scratching the surface? By addressing these questions, we aim to provide a data-driven reassessment of microbial diversity in soil and to establish a benchmark for future metagenomic studies of complex ecosystems.

## Methods

### Sample collection, DNA extraction, and metagenomic sequencing

The A horizon of the soil type Cambisol was sampled from the Schönbuch Forest, near Tübingen, Germany (the same site previously reported by [19]), on 31 May 2022. High molecular weight metagenomic DNA was isolated using a protocol described in detail in our previous studies [19, 20]. Genomic integrity was assessed using pulse-field capillary electrophoresis with the Genomic DNA 165 kb Analysis Kit on a FemtoPulse (Agilent) instrument. Quantitation of DNA was assessed using the dsDNA High Sensitivity Assay on a Qubit 3 fluorometer (Thermo Fisher), and purity was assessed by Nanodrop. A total of 2.4 µg of genomic DNA was used as input for the library preparation with the 1D Ligation Kit SQK-LSK109-XL Sequencing Kit (ONT). Four PromethION R9 flow cells were utilised for long-read Nanopore sequencing, each loaded with 600 ng (50 fmol) of genomic DNA. The raw signal data from the PromethION runs were basecalled with guppy (v 5.0.7) in high-accuracy mode. For complementary short-read sequencing, three NovaSeq 6000 flow cells were loaded with the same isolated DNA; two of them ran with 200 cycles, and one with 300 cycles.

A total of 148 Gbp of Nanopore sequencing data was generated, with a read length N50 of 12.2 kbp. The three Illumina sequencing runs yielded a total of 122 Gbp raw data, with a mean Q-score of 35. The raw Illumina reads were adapter and quality trimmed using fastp [21] (v 0.23.4) with default settings.

### Metagenomic assembly

Nanopore reads from all four runs were pooled together and assembled using metaFlye (v2.9.5-b1801) [22] with the `--meta` option optimised for metagenomic data, and the `--nano-raw` option for the error-prone Nanopore reads. The resulting draft assembly was polished with one round of medaka [23] (v 2.0.1) using the `r941_prom_hac_g507 model` and the `--bacteria` flag. The medaka polished assembly was further corrected using the trimmed Illumina reads in a single round of NextPolish [24] (v 1.4.1) polishing, with parameters `-max_depth 100` for short-read mapping with bwa [25] (v 0.7.18), and `-min_read_len 1k -max_depth 100 -x map-ont` for long-read mapping with minimap2 [26] (v 2.28-r1209).

In parallel, an Illumina-only assembly was performed using MEGAHIT [27] (v1.2.9) with default metagenomic assembly parameters. All subsequent analyses were based on the hybrid Nanopore-Illumina assembly, which exhibited superior contiguity (contig N50 = 77.8 kbp) and total assembled length (10.5 Gbp).

### Taxonomic analysis

Raw reads were taxonomically classified using Metabuli [28] (v 1.0.8), against the precomputed database provided by the authors, which includes GTDB release 214.1 [29] and the human T2T genome [30]. Classification was performed with parameters `--seq-mode 2` for Illumina reads, and `--seq-mode 3` for Nanopore reads. Contigs from the final hybrid assembly were taxonomically classified using MMSeqs2 [31] (v 16.747c6) using the `easy-taxonomy` module, with GTDB release 214.1 [29]. A contig was classified as “cultured” at rank *R*, if its assignment at *R* belongs to a taxon that has at least one genome in GTDB originating from an isolate sequencing; as “uncultured” if an assignment exists at rank *R*, but all representatives of the taxon come from MAG/SAG sequencing efforts; and as “unclassified” if there is no assignment at rank *R* due to insufficient similarity or conflicting similarities. The information on assigned genomes belonging to a cultured or uncultured representative was retrieved from GTDB metadata files.

### Rarefaction analysis

Microbial diversity and sequencing coverage were assessed using multiple complementary approaches. K-mer frequency analysis was conducted with ntCard [32] using 15-mers and 17-mers for both Nanopore and Illumina datasets. Nonpareil [33] (v3.3.3) was used to estimate sequence coverage and to project additional sequencing efforts required for complete community recovery using the pooled set of Illumina reads. Full-length 16S rRNA gene sequences were extracted from Nanopore reads using Barrnap [34] (v0.9) (sequences flagged as “partial” were excluded). We performed rarefaction by subsampling them at 10% increments (10-100%), with three independent replicates per depth (distinct random seeds). For each subset, we clustered the subsampled sequences with VSEARCH [35] (v2.15.2), reporting both exact uniques (100% dereplication) and the number of representatives at fixed identity thresholds (e.g. 97%, 94%, 88%, 85%, 70%, 61%). VSEARCH was run with `--strand both --iddef 1 --qmask none` and `--cluster_size` options for maximum sensitivity. In addition, conserved single-copy marker genes from the bac120 set [13] were identified in Nanopore reads using the LAST aligner (v 1615) in frameshift mode (-F 15) against the GTDB bac120 reference set, and quantified to independently assess the taxonomic diversity.

### Functional annotation, clustering, and biosynthetic gene cluster discovery

The final polished assemblywas annotated using Bakta [36] (v 1.11.0, database version 5.1). The identified coding sequences (CDS) were further annotated for their COG categories [37], eggNOG orthologous groups [38], KEGG ontologies, pathways, modules, reactions [39], Gene Ontologies [40], CAZy enzymes [41], and Pfam domains [42], using eggNOG-mapper [43] (v2.1.13) (Supplementary Figure S1 and the Zenodo repository).

We also clustered these predicted CDS with MMseqs2 linclust [44] (v18.8cc5c) across identity thresholds of 90%, 80%, 70%, 60%, and 50%. Clustering used an 80% bidirectional coverage requirement (`cov-mode 1, -c 0.8`), low-complexity masking, and the fast greedy set-cover algorithm. For each threshold, MMseqs2 produced representative-member assignments; we then performed bootstrap rarefaction on the member-representative mapping by repeatedly subsampling CDSs to increasing depths and counting the number of unique clusters observed (100 steps, and 250 bootstraps). Replicates were aggregated as mean *±* confidence intervals to generate rarefaction curves of unique protein clusters against the number of sampled CDSs.

Biosynthetic gene clusters (BGCs) were identified using antiSMASH [45] (v7) on the entire hybrid assembly, with the options `--fullhmmer --clusterhmmer --tigrfam --asf --cc-mibig --cb-general --cb-subclusters --cb-knownclusters --pfam2go --rre --smcog-trees --tfb`. Identified BGCs from the hybrid assembly were subsequently clustered into gene cluster families (GCFs) using BiG-SCAPE [46] (v2 beta5), based on domain architecture and sequence similarity, at a category-level distance threshold of 0.2. The novelty of BGCs was assessed by comparisons against the MIBiG 4.0 [47] database by the BiG-SCAPE search; and independently against the BGC Atlas [48] database by using its built-in BiG-SLiCE [49] search functionality at the pre-defined distance threshold of 0.4. The BGCs were classified as “novel” when they did not cluster with any MIBiG reference BGC or BGC Atlas reference GCF.

### Reconstruction of metagenome-assembled genomes

Metagenome-assembled genomes (MAGs) were recovered using a combination of complementary binning strategies to maximise genome quality. Binning was first performed independently on the final NextPolish-corrected assembly using COMEBin [50], MetaDecoder [51], SemiBin2 [52], and VAMB [53], executed in a CUDA-enabled environment when possible.

COMEBin (v 1.0.4), together with CheckM (v 1.1.3) [54], was run on alignments generated by minimap2 [26] (v2.28-r1209), with the following parameters: `NUM_VIEWS=6, TEMPERATURE=0.07, EMBEDDING_SIZE=2048, COVERAGE_EMBEDDING_SIZE=2048`, and `BATCH _SIZE=1024`. MetaDecoder (v 1.1.0) was executed with default parameters according to the authors’ recommendations. SemiBin2 (v 2.1.0) was run in both `--self-supervised` and `--semi-supervised` modes, specifying soil as the environment type and long_reads as the sequencing type. Gene prediction was performed using Prodigal (v 2.6.3) [55], and the GTDB v95 database [29] was used alongside the previously described minimap2 mappings of the long reads.

For VAMB (v 4.1.4) [53], six binning results were generated by combining default and taxonomy-aware binning modes with three input configurations: long reads, short reads, and both. Abundance profiles were computed using Strobealign [56] (v 0.15.0). For Tax-VAMB [57], contig-level taxonomic annotations were derived using Metabuli [28] against the GTDB reference database [29].

In order to produce a final, unified metagenome-assembled-genome binning, the binning results from the above-mentioned ten runs were collected, and refined using MAGScoT [58] (v 1.1). MAGScot was run on gene-calling results from prodigal [55] (v 2.6.3), and hmmsearch results from the HMMER package [59] (v 3.4), using Pfam [42] and TIGRFAMs [60] HMM profile databases.

The resulting MAGs were evaluated for their completeness and contamination with CheckM2 [61] (v 1.0.2). Pairwise genomic distances were computed using Mash [62] (v 2.3) with default parameters. Taxonomic assignments were performed with GTDB-Tk [63] (v 2.4.0) using the GTDB r220 release [29] in classify mode. A phylogenetic tree of high- and medium-quality MAGs was constructed with GTDB-Tk in *de novo* mode and visualised with iTOL [64].

These MAGs were also annotated using Bakta [36] and antiSMASH [45] as described above, both to establish links between BGCs and MAGs and to provide a reference annotation set for future studies (available in the Zenodo repository). Our hybrid (ONT+Illumina) analysis was executed via a Nextflow DSL2 pipeline released on WorkflowHub [65].

## Results

### Ultra-deep hybrid sequencing and assembly of a temperate forest sample

To evaluate the limits of current sequencing technologies in capturing soil microbial diversity, we generated an ultra-deep metagenomic dataset from a single sample collected in the Schönbuch forest, a temperate mixed forest in southwestern Germany. The sample was taken from the A horizon of a Cambisol soil, a type of soil known for its rich microbial diversity and favourable characteristics for DNA extraction. This particular site and soil type were previously shown to harbour high biosynthetic potential and taxonomic richness [19], and were therefore selected as a representative model system for deep metagenomic exploration.

We extracted total environmental DNA from this soil and sequenced it using a hybrid strategy that combined 148 Gbp of ONT long-read and 122 Gbp of Illumina short-read sequencing. Nanopore reads had a read length N50 of 12.2 kbp, while Illumina reads achieved a mean Q-score of 35. Together, this 270 Gbp dataset represents one of the deepest single-sample soil metagenomes reported to date, and provides a unique opportunity to empirically assess taxonomic and functional complexity.

We employed a hybrid assembly strategy to combine the complementary strengths of long and short reads. Long-read Nanopore data were assembled and subsequently polished using Illumina data, resulting in 10.5 Gbp of assembled sequence with a contig N50 of 77.8 kbp. For comparison, an Illumina-only assembly produced a more fragmented assembly with an N50 of only 865 bp and a total length of 9.2 Gbp, confirming the advantages of long-read inclusion.

### Taxonomic analysis reveals unprecedented levels of novelty

Taxonomic classification of both the raw reads and the assembled contigs revealed a highly diverse and complex microbial community, comprising members from 69 different phyla. As expected for soil environments, the community was dominated by taxa such as Pseudomonadota and Acidobacteriota (Figure 1A), along with representatives from many other phyla. A small fraction (approximately 1%) of the reads originated from eukaryotes, while 8% remained unclassified, indicating the presence of deeply novel sequences absent from current databases.

**Figure 1.**
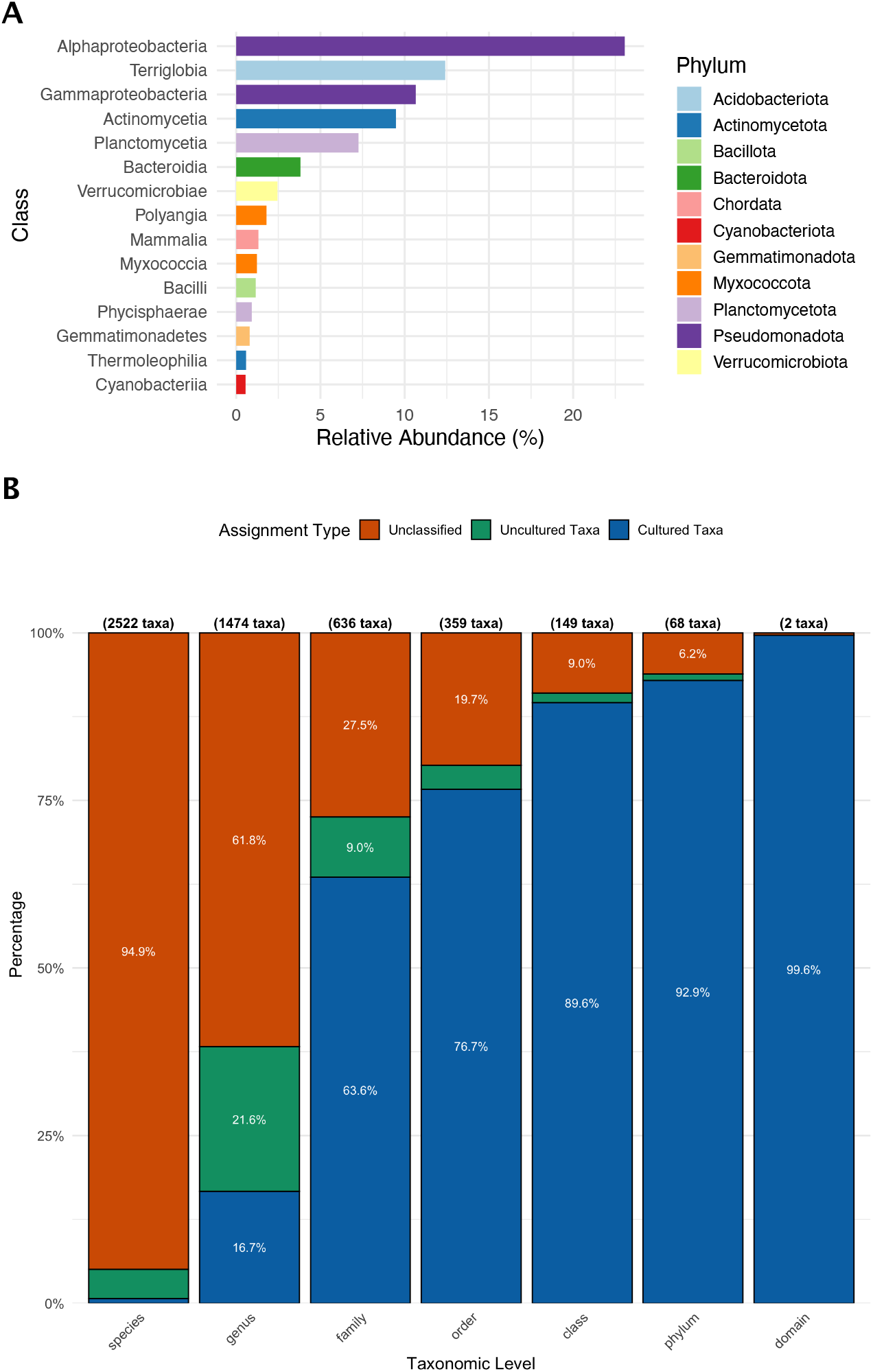
Taxonomic summary of the assembled contigs. **(A)** Class-level relative abundances of the assembled contigs, coloured by the phyla they belong to. **(B)** Taxonomic novelty levels of the assembled contigs across different ranks. Bars show the proportion of total assembled bases assigned to: cultured taxa (blue), where at least one GTDB representative is an isolate genome; uncultured taxa (green), where all GTDB representatives are MAGs/SAGs only; or unclassified (orange), where no assignment was returned at rank *R* (e.g. insufficient similarity or conflicting hits). The number of taxa identified at each level is shown above each bar. Most sequences could only be classified at higher taxonomic ranks, while species- and genus-level assignments were predominantly unclassified.

Taxonomic analysis of the assembled contigs (Figure 1B) further demonstrated that a substantial portion of the dataset originates from uncultivated organisms across all taxonomic ranks. While soil has long been recognised as a reservoir of uncultivated diversity, with historical estimates suggesting 90–99% of taxa remain uncultured, our data now provide a quantitative confirmation of this.

At the species level, more than 99% of the assembled contigs could not be assigned to any taxon with a cultured representative. Only 0.7% of contigs could be linked to species with at least one cultured strain, and 16.7% to a genus with cultured members. At the species level 4.5%, and at the genus level 21.6% of the contigs were linked to a previously sequenced MAG/SAG; however, they did not have a cultured representative. 94.4% and 61.8% of the assembled data could not be assigned to any genome present in the databases at species and genus levels, respectively.

Even at broader taxonomic levels, substantial novelty is evident: 9% of contigs could not be assigned to any known class (cultured or uncultured), and 6.2% could not be linked to any known phylum (Figure 1B). Among the contigs that were classified, a substantial fraction was affiliated with taxa represented exclusively by uncultivated lineages, known only from metagenomic or single-cell sequencing studies. These patterns are in contrast to metagenomic datasets from more tractable environments, such as the human gut or ocean microbiomes, where a larger fraction of sequenced data can be linked to cultivated taxa [66, 67].

### Soil diversity remains undersampled

To evaluate whether ultra-deep sequencing could fully capture the microbial diversity present in the sample, we applied a combination of k-mer analysis, rarefaction based on marker genes, full-length 16S rRNA genes, and predicted CDSs. All approaches consistently indicated that, despite the unprecedented depth of sequencing, the dataset remains far from saturating the underlying biological diversity.

K-mer analysis, which examines the frequency distribution of short subsequences within the sequencing reads, showed a distribution heavily weighted toward low-frequency k-mers (Figure 2A). Most k-mers occurred only once or a few times, reflecting the high prevalence of unique or rare sequences. This pattern is characteristic of highly diverse communities and indicates the presence of many low-abundance taxa whose genomes were either incompletely covered or missed entirely in assembly. A long right-hand tail, corresponding to high-frequency k-mers, likely originated from abundant taxa or repetitive genomic elements (such as rRNA genes). These patterns were consistent across Nanopore and Illumina datasets and highlight the extreme sequence heterogeneity of the sample (Figure 2A, and Supplementary Figures S2 and S3).

**Figure 2.**
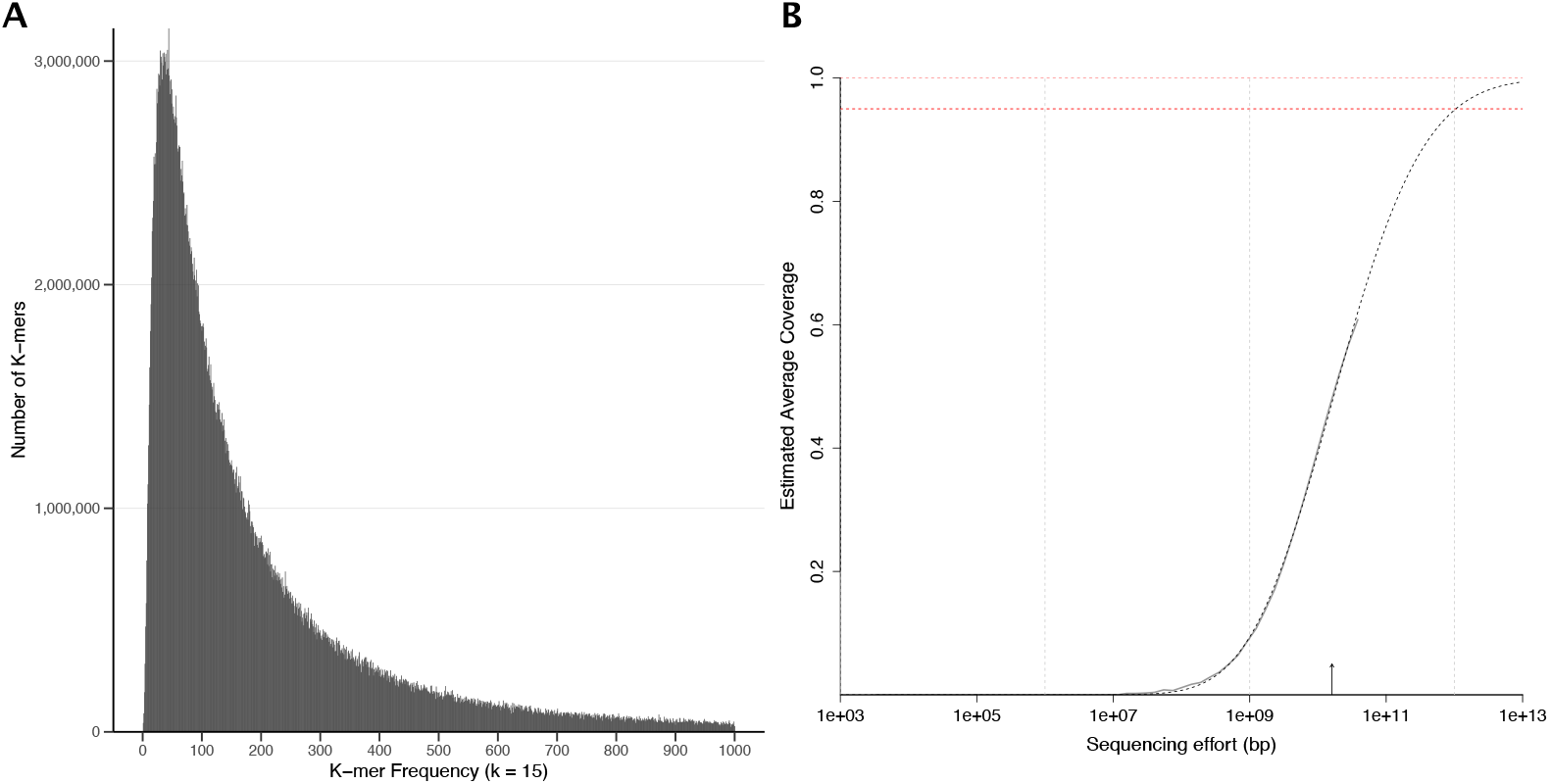
Sequencing depth diagnostics show that ultra-deep sequencing still undersamples the total soil diversity. **(A)** 15-mer spectrum of Nanopore reads from the soil metagenome. Bars show the number of distinct 15-mers observed at each copy number. The pronounced peak at low frequencies reflects a very large pool of unique or low-coverage k-mers, indicating extreme sequence diversity. The long right-hand tail originates from higher coverage k-mers, likely derived from abundant taxa and repetitive genomic regions. **(B)** Nonpareil analysis estimating the sequencing coverage of the pooled Illumina dataset. The curve shows the relationship between sequencing effort and estimated coverage, with the solid line representing the observed dataset and the dashed extrapolation indicating expected gains with additional sequencing. The analysis suggests that substantially more sequencing effort would be required to achieve near-complete coverage of the community.

Rarefaction analysis with Nonpareil further supported this conclusion. Based on read redundancy, the model projected that approximately 10 terabases of sequencing data would be required to achieve near-complete coverage of the microbial diversity in this single soil sample; almost 50 times more than our current sequencing effort (Figure 2B). This analysis confirms that even the most comprehensive sequencing efforts to date fall short of fully resolving the taxonomic complexity of soil microbiomes.

In parallel, we extracted a total of 19 738 full-length 16S rRNA gene sequences from Nanopore reads (Supplementary Figure S4) and clustered them at various sequence identity thresholds. At 100% and 97% identity, the number of observed 16S rRNA gene clusters continued to increase linearly with sequencing depth, showing no indication of saturation (Figure 3A). A complementary analysis based on 120 conserved single-copy bacterial marker genes showed similar trends, with between 9000 and 59 000 unique copies detected per gene across the dataset (Supplementary Figure S5). While at lower clustering thresholds, the number of representative 16S rRNA gene sequences began to plateau; at higher identity thresholds, the curves remained unsaturated. Similarly, the rarefaction of clustered CDSs showed that observed richness continued to increase even after sampling 10 million CDSs, irrespective of the clustering threshold applied. These steep, non-saturating curves indicate not only unresolved taxonomic diversity but also a remarkably deep functional repertoire present in the soil community.

**Figure 3.**
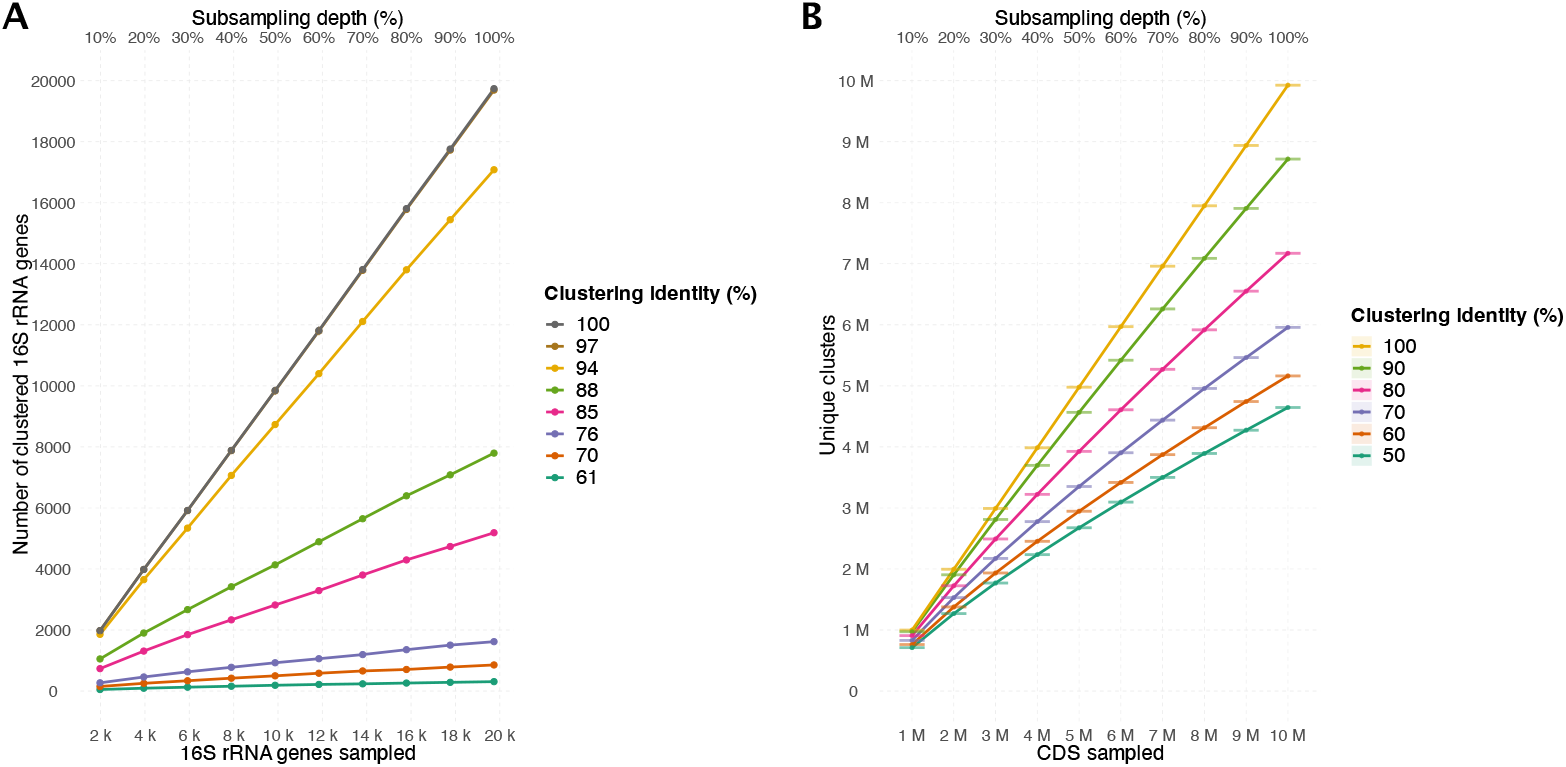
Rarefaction curves of full-length 16S rRNA gene sequences extracted from Nanopore reads (A) and CDSs (B) at different clustering thresholds. **(A)** The number of observed full-length 16S rRNA gene sequences is plotted against the number of sampled genes at clustering thresholds ranging from 61% to 100% sequence identity. The curves at high-identity thresholds, such as 97% and 100%, increase in a near-linear fashion, indicating that the vast majority of taxonomic diversity remains undiscovered, while the increase is less steep at lower clustering thresholds. **(B)** Rarefaction curves of predicted coding sequences (CDSs) from the final assembly, clustered at identity thresholds from 50% to 100%, with a bi-directional coverage requirement of 80%. The number of unique protein clusters continues to rise steeply even after sampling 10 million CDSs, demonstrating that the functional gene repertoire of the soil community is also far from saturated.

Taken together, these results provide robust and multi-dimensional evidence that the soil microbial community remains vastly undersampled, even with sequencing efforts far beyond typical practice. The failure to reach saturation was especially apparent at the species level, where the rarefaction curves continued to rise linearly across all subsampling depths. This indicates an exceptional degree of fine-scale taxonomic resolution, with many closely related but genomically distinct lineages present at low abundance. Similarly, functional clustering of coding sequences revealed a parallel pattern: at high identity thresholds, the number of distinct CDS clusters continued to grow steeply with sequencing depth, while at lower thresholds, sequences clustered more readily, reflecting both the breadth of novel gene diversity and underlying functional redundancy across lineages.

Such strain- and species-level diversity is particularly important given that many ecologically and functionally relevant traits, such as niche specialisation, symbiosis, and secondary metabolite biosynthesis, can differ even between closely related strains. In more constrained environments, such as the human gut, species-level diversity tends to plateau after a few hundred genomes [68], and marine planktonic communities exhibit relatively stable core microbiomes [69]. By contrast, the soil microbiome appears to exhibit near-limitless diversity at the genomic level.

These findings not only demonstrate the limitations of current sequencing depths but also call into question the accuracy of historical species richness estimates derived from much shallower data. They suggest that achieving anything approaching species-level saturation in soil will likely require terabase-to petabase-scale sequencing efforts, combined with refined computational methods to resolve low-abundance and highly diverse taxa. They demonstrate that many of the diversity estimates historically used in soil microbiome research, often based on much shallower sequencing, likely underestimate the true scale of taxonomic complexity.

### Long-read metagenomics recovers thousands of novel and genome-linked BGCs

From the hybrid metagenomic assembly, we identified a total of 11 381 BGCs, including 5652 classified as complete, reflecting the improved contiguity made possible through long-read sequencing.

This proportion of complete clusters (∼50%) represents a substantial improvement over previous metagenomic surveys, where fewer than 10% of BGCs are typically annotated as complete [48].

Clustering the identified BGCs with BiG-SCAPE resulted in 10 215 gene cluster families, indicating that most BGCs in our dataset are non-redundant and functionally distinct. The size distribution of GCFs (Figure 4A) was highly skewed: most families consisted of singletons or doubletons, with only a handful of larger families, further illustrating the functional uniqueness and fine-grained metabolic specialisation within the community. The largest GCF contains 115 ribosomally synthesised and post-transcriptionally modified peptide (RiPP) BGCs. The cumulative ordering of the GCFs by size shows that the largest eight families account for 3.1% of all BGCs, whereas each of the remaining over ten thousand families contributes only very little to the total number (Figure 4B).

**Figure 4.**
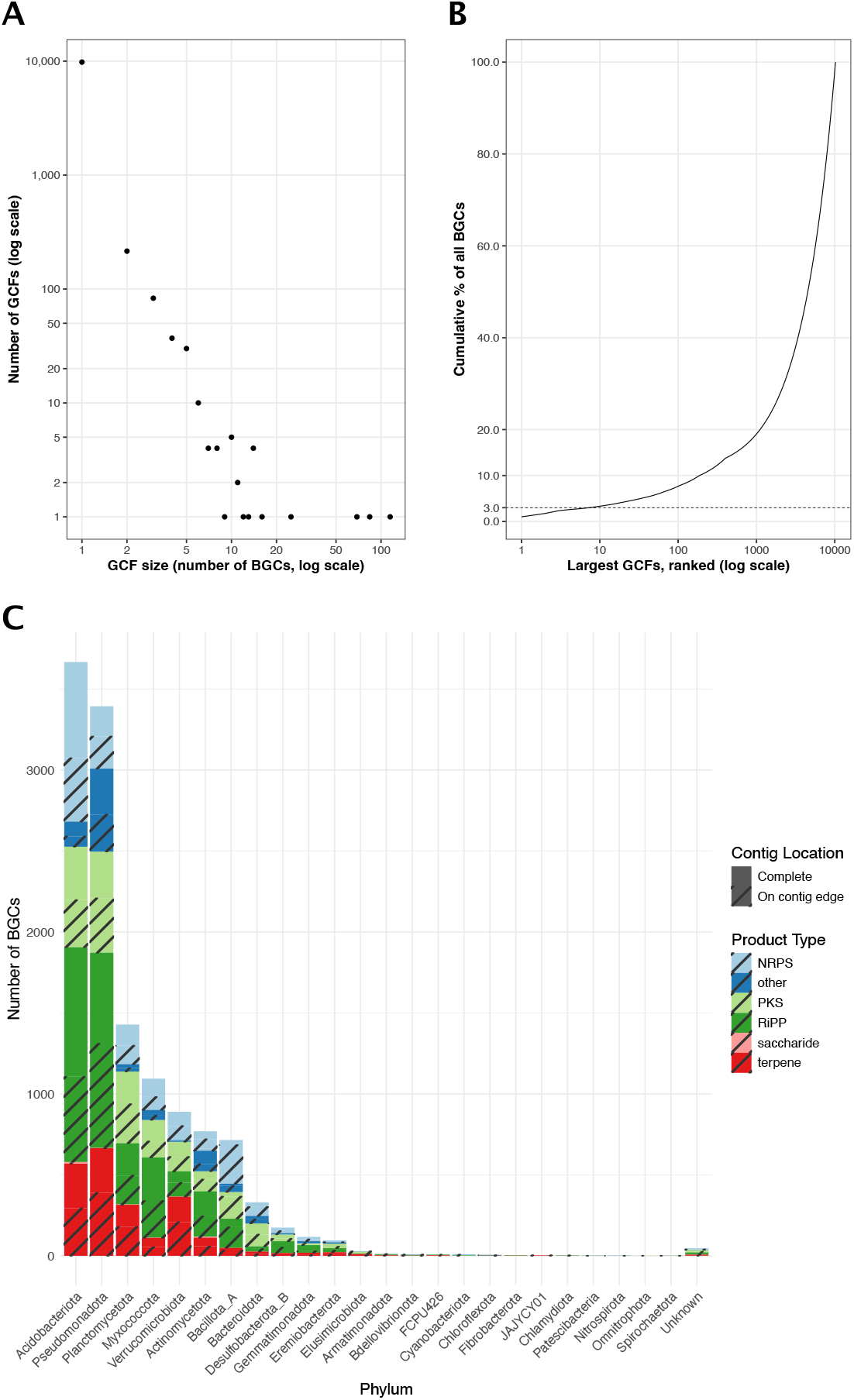
The diversity and taxonomic composition of the biosynthetic gene clusters. **(A)** The size distribution of gene cluster families (GCFs). Each dot represents one distinct GCF size in the dataset (11 381 biosynthetic gene clusters grouped into 10 215 GCFs). The x-axis shows the family size (number of BGCs), the y-axis shows the number of families of that size; both axes are log_10_ -scaled. The plot reveals a heavy tail: 9813 families (96.1 %) are singletons, only 20 families contain ≥10 BGCs, and the largest family comprises 115 BGCs. **(B)** Cumulative contribution of the largest GCFs to the total BGC dataset. Families are ranked from largest to smallest along the log_10_ -scaled x-axis; the y-axis tracks the running percentage of all 11 381 BGCs accounted for as each additional family is added. The eight largest GCFs explain 3.1% of all BGCs, whereas 7939 families are required to reach 80%. The rapid rise followed by a very shallow tail shows that a small handful of families contribute to about 3% of BGCs in the dataset, while thousands of minor families are unique. **(C)** Distribution of biosynthetic gene cluster (BGC) product types across bacterial phyla. Each bar represents the total number of BGCs assigned to a given phylum, coloured by predicted product type. Striped regions indicate the proportion of BGCs classified as “on-contig-edge” by antiSMASH. Phyla are ordered by total BGC count, with “Unknown” denoting contigs for which no confident phylum-level classification could be assigned.

Taxonomic classification of the BGC-containing contigs revealed that biosynthetic potential is broadly distributed across the phylogenetic spectrum. While well-known producers such as Actinomycetota and Pseudomonadota were represented, we also detected numerous BGCs in less-studied phyla such as Verrucomi-crobiota and Acidobacteriota (Figure 4C). These groups have historically been overlooked due to cultivation barriers but appear to harbour substantial secondary metabolic capacity.

Only 108 BGCs (∼ 1%) had significant matches to known GCFs in the BGC Atlas database, which compiles nearly two million BGCs from metagenomic sources, and none clustered together with reference BGCs from MIBiG. This confirms earlier observations that soil metagenomes are rich in biosynthetic novelty [15, 70], but also demonstrates that even with massive reference expansions, a typical soil still contains an overwhelming majority of uncharacterised BGCs. Notably, our dataset contains more novel BGCs than reported in most large-scale studies of marine or human microbiomes, where the rate of novel BGC discovery is often constrained by lower microbial diversity and shorter contigs [70, 48].

### Hybrid assembly recovers hundreds of genomes with high taxonomic and biosynthetic novelty

From the hybrid assembly, we recovered 837 metagenome-assembled genomes (MAGs), of which 466 met the MIMAG standards for high- or medium-quality [71]. High- and medium-quality MAGs ranged in size from 1 Mbp (Patescibacteria) to 15 Mbp (Planctomycetota) (Figure 5 and Supplementary Table S1). All MAGs recovered represented unique species-level units, defined by > 0.05 pairwise Mash distance (≥ 95% ANI). These genomes spanned a wide range of bacterial phyla typical of soils, including Acidobacteriota, Pseudomonadota, Verrucomicrobiota, Actinomycetota, and Myxococcota. Remarkably, only four of the 837 MAGs had species-level matches (≥95% ANI) in the GTDB database, and none had cultured representatives.

**Figure 5.**
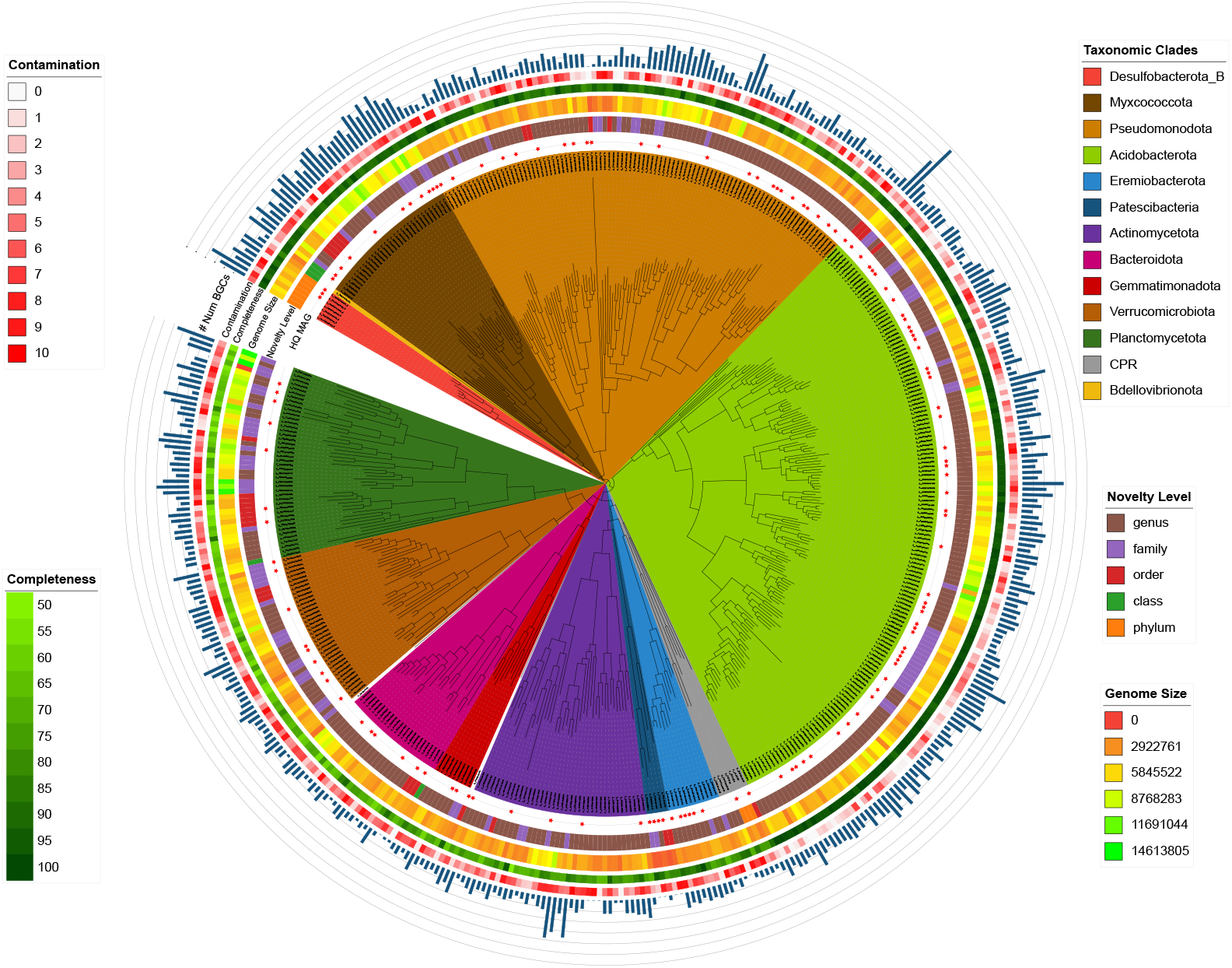
Genome quality statistics and taxonomic classification of newly reconstructed high-quality and medium-quality metagenome-assembled genomes (MAGs). Each leaf represents a MAG, coloured by its assigned phylum. Concentric rings around the leaves indicate, from innermost to outermost: (i) HQ MAGs, (ii) novelty level (genus to phylum), (iii) genome size (bp), (iv) completeness (%), (v) contamination (%), and (vi) the number of BGCs encoded by the MAG (range: 0-30).

Thirteen MAGs were assigned to phyla with no cultured members, such as Desulfobacterota, and candidate phyla radiation (CPR) groups FCPU426 and JAJYCY01, harbouring between 2 and 19 BGCs (Figure 5). At other taxonomic levels, 9, 45, 155, and 611 MAGs belong to novel lineages that correspond to class, order, family, and genus levels, respectively.

Many of the recovered MAGs are assigned to taxonomic groups that do not correspond to any known families or even classes in current reference databases (Figure 5). This points to a substantial fraction of microbial diversity in soil that remains undescribed, even at higher taxonomic ranks. In addition to their taxonomic novelty, many of these genomes contain a large number of BGCs, in some cases reaching 30 distinct clusters within a single genome. This suggests that the potential for secondary metabolite production is not restricted to well-characterised soil taxa (such as Actinomycetota), but is also widespread among poorly characterised lineages.

Of all BGCs identified, 6279 could be linked to metagenome-assembled genomes, enabling genome-resolved exploration of natural product potential. This represents one of the largest genome-resolved inventories of BGCs from a single environmental sample to date.

The abilityto recover thousands of complete and genome-linked BGCs from a single soil metagenome demonstrates the power of ultra-deep long-read sequencing in capturing microbial metabolic potential. In addition, the high novelty rate and wide taxonomic distribution of these clusters emphasise a vast and largely untapped chemical repertoire unlikely to be captured through shallow sequencing or cultivation-based surveys alone.

## Discussion

### Soil diversity still outruns ultra-deep sequencing

Our study presents one of the most comprehensive metagenomic investigations of a single soil microbiome to date, combining ultra-deep Oxford Nanopore and Illumina sequencing to yield nearly 300 Gbp of data. This effort enabled high-quality hybrid assemblies and genome-resolved analysis of taxonomic and functional potential in a temperate forest soil sample, providing an empirical benchmark for the capabilities and current limitations of metagenomic sequencing in one of the most complex microbial habitats on Earth.

Despite the ultra-deep sequencing depth, our analyses demonstrate that the microbial diversity of the sample remains far from saturated. Rarefaction curves, k-mer profiles, 16S rRNA gene and marker gene analyses, and functional clustering of CDSs consistently indicate that substantially more sequencing would be required to approach near-complete coverage of the community diversity, with a non-parametric model estimate on the order of ∼ 10 Tbp of raw sequencing data to reach high-coverage targets (Figure 2). By comparison, MAG recovery in human-gut communities reaches saturation with ∼ 5–10 Gbp per sample, and 25 Gbp is estimated to be enough even to classify taxa of very low abundances (< 1e-06) [72]. Similarly, Ni et al. [73] estimate this number to be around 7 Gbp to detect gene content of organisms with relative abundances over 1% in human faecal samples.

Ultra-deep campaigns such as Tara Oceans have shown that planktonic marine communities also benefit from multi-terabase sequencing, yielding thousands of novel genes and genomes and greatly refining oceanic reference catalogues. Yet, the incremental discovery of truly new sequence space in the ocean has begun to flatten at these depths [69, 74, 75]. In soils, by contrast, every additional sequencing effort still brings a steeper growth of previously unseen sequence diversity, and MAGs, with no hint of saturation.

Because our analysis derives from a single temperate Cambisol soil sample, these results should be read as a depth benchmark rather than a census of soils at a global scale. We expect the heavy-tailed diversity and lack of saturation to generalise to other complex soils; however, the taxonomic composition and biosynthetic potential may vary with soil type, season, agricultural use, and other factors.

During the revision of this manuscript, three independent studies using broader sampling and alternative designs reached similar conclusions, placing our single-site depth benchmark in a broader context. A multi-habitat long-read resource sequenced 154 Danish soils with an emphasis on expanding reference space across habitats, and recovered more than 15 000 previously undescribed species and thousands of complete BGCs, demonstrating how long reads broadly improve genome recovery and downstream annotation across terrestrial microbiomes [76]. They also demonstrated a near-linear rarefaction relationship between the number of MAGs that are recovered from the soil samples with the number of species they correspond to. Complementarily, another analysis of publicly available short-read sequencing efforts of soil samples estimated that multi-terabase sequencing efforts per sample may be needed to approach near-complete saturation, consistent with our observation that soil diversity remains unsaturated at current depths [77]. They also demonstrate that large co-assembly efforts of short-read soil metagenomics improve the coverage, yet still fall short of capturing it completely. Additionally, Burian et. al. [78] introduced the first terabase-scale long-read sequencing effort of a single soil sample, focusing on the recovery of nonribosomal peptide (NRPS) BGCs. They generated 2.5 Tbp of raw long-read data from one forest soil, where the NRPS discovery scaled linearly up to the 2.5 of the generated data, not capturing the full diversity of NRPS BGCs in the single soil sample.

### Uncharted novelty of soil microbes and their functional capabilities

Taxonomic profiling of assembled contigs revealed the presence of a diverse range of phyla in the soil, including the expected dominance of well-known soil phyla, such as Acidobacteriota, Actinomycetota, and Pseudomonadota, alongside under-studied lineages. Yet, over 95% of contigs could not be assigned to a species currently present in databases, and less than 0.7% could be linked to a species with at least one cultured representative. We recovered MAGs that represent candidate novel phyla, highlighting the ability of genome-resolved metagenomics to reveal deeply divergent lineages missed by conventional approaches. Only four MAGs recovered in this study share ≥ 95% ANI with genomes in GTDB, and none correspond to cultured isolates, reaffirming how incompletely current databases represent soil ecosystems. These lineages not only expand the bacterial tree of life but, in many cases, also harbour unique biosynthetic gene clusters, suggesting functional capacities that remain entirely uncharted.

The scale of biosynthetic diversity uncovered is similarly striking. We identified over 11 000 BGCs, more than 5600 of which are complete and many of which could be linked to MAGs. Fewer than 1% of these BGCs matched known families in public databases, suggesting that soils remain a massive and underexplored reservoir for natural product discovery. In other words, the chemistry encoded in a single forest soil sample still sits almost entirely outside the reach of current reference catalogues. The heavy-tailed GCF distribution we observe, where more than 96% of families are singletons, suggests that every additional increment of sequencing continues to reveal functionally distinct BGCs, rather than finding copies of known ones.

These BGCs come from virtually every major bacterial phylum detected in the sample, from well-studied Actinomycetota and Pseudomonadota to under-explored Acidobacteriota, Verrucomicrobiota, and candidate phyla radiation groups. This phylogenetic breadth and the underestimated functional diversity of soil microbes indicate that the soil’s metabolic repertoire is not restricted to a few prolific lineages but is dispersed across the microbial tree of life, with implications for microbial interactions, nutrient cycling, defensive traits, and overall ecosystem stability.

Furthermore, our dataset shows that long-read sequencing can address common limitations of metagenomics approaches, including incomplete recovery of BGCs and fragmented assemblies, thereby enabling genome-resolved assessments of secondary metabolism. In contrast to earlier studies depending exclusively on short-read data [48], our approach yields a high proportion of complete BGCs and facilitates the association of biosynthetic capacity with uncultured taxa.

### Conclusions

Together, these results redefine the scale of the unknown in soil microbiomes. They demonstrate that even the deepest sequencing efforts to date barely scratch the surface of microbial and metabolic diversity, particularly in complex environments. Moving forward, terabase-to petabase-scale sequencing will likely be necessary to achieve species-level saturation in soil. However, deeper sequencing alone will not be sufficient. Advances in bioinformatics for assembly, binning, and functional annotation, combined with complementary data from metatranscriptomics, metabolomics, and single-cell approaches, will be essential for translating sequence data into ecological and biochemical insight.

In conclusion, this study sets a new reference point for what is currently achievable in soil metagenomics. By combining ultra-deep sequencing with advanced computational methods, we provide an updated, data-driven view of soil microbial and biosynthetic diversity. The dataset offers an empirical baseline for evaluating sequencing depth, genome recovery, and biosynthetic novelty in complex environments. As such, it can serve as a valuable resource for guiding future studies, benchmarking new tools, and refining diversity estimates. Our findings highlight both the power of current technologies and the need for continued investment in sequencing and analysis to uncover the vast, largely uncharacterised microbial life in soil ecosystems.

### Potential implications

Beyond its soil-ecological focus, this ultra-deep, genome-resolved dataset can (i) act as a benchmark for soil diversity in future studies. (ii) It supplies hundreds of genomes from previously unseen lineages to extend genome databases and taxonomies such as MGnify [79, 80] and GTDB [29], and refine the bacterial tree of life. (iii) It contains >11 000 largely novel and complete BGCs, many linked to host genomes, which can extend the secondary metabolite databases such as BGC Atlas [48] and the Secondary Metabolism Collaboratory (SMC) [81]; (iv) and this phylogenetically diverse and novel collection of genomes and BGCs can be used as a training set for machine-learning models that predict compound functions [82] and structures [83], and the ecological traits of microbes [84].

## Availability of source code and requirements

- Project name: Soil Metagenome Pipeline
- Project home page: soil-metagenome-pipeline https://github.com/ZiemertLab/
- WorkflowHub SEEK ID: https://workflowhub.eu/workflows/1960?version=1
- License: GNU General Public License v3.0

System requirements

- Operating system(s): Linux
- Programming language: Nextflow DSL2, Python, bash
- Other requirements: conda, SLURM
- Hardware requirements: >64 threads, >128GB RAM

## Data availability

The raw sequencing data generated during the current study are available in the ENA repository [1]. The identified biosynthetic gene clusters (BGCs), and extracted MAGs are available via Zenodo for unrestricted reuse [2, 3]. The computational analysis pipeline has been made available on GitHub [85] with the GNU General Public License v3.0 and deposited on workflowhub.eu [65] as a NextFlow pipeline.

## Additional Files

*Supplementary Fig. S1*. Distribution of predicted proteins into COG categories.

*Supplementary Fig. S2*. 17-mer spectrum of Illumina reads.

*Supplementary Fig. S3*. 17-mer spectrum of Nanopore reads

*Supplementary Fig. S4*. Length distribution of the 16S rRNA gene sequences extracted from raw Nanopore reads.

*Supplementary Fig. S5*. The number of occurrences of each marker gene from bac120 set in raw Nanopore reads.

*Supplementary Table S1*. The details of the extracted MAGs, including their completeness and contamination values, taxonomic classification, assembly statistics, genome annotations statistics, and the number of BGCs identified.

## Declarations

## List of abbreviations

ANI: average nucleotide identity
BGC: biosynthetic gene cluster
bp: base pairs
CDS: coding sequence
CPR: Candidate phyla radiation
DNA: deoxyribonucleic acid
ENA: European Nucleotide Archive
Gbp: giga (billion) base pairs
GCF: gene cluster family
GTDB: Genome Taxonomy Database
HQ: high quality
MAG: metagenome-assembled genome
MIBiG: Minimum Information about a Biosynthetic Gene Cluster
MIMAG: Minimum Information about a Metagenome-Assembled Genome
MQ: medium quality
NGS: next-generation sequencing
NRPS: non-ribosomal peptide synthetase
kbp: kilo (thousand) base pairs
Mbp: mega (million) base pairs
ONT: Oxford Nanopore Technologies
PKS: polyketide synthase
Q-score: quality score (Phred)
RiPP: ribosomally synthesised and post-transcriptionally modified peptide

## Ethical Approval

Not applicable

## Consent for publication

Not applicable

## Authors’ Contributions

C.B. and N.Z. wrote the main manuscript. C.B. performed the data analysis and prepared the figures. T.N. conducted the soil sampling and DNA isolation. E.A.B. carried out the Nanopore sequencing. C.G. performed basecalling and preliminary quality control of the sequencing data. C.B., S.O., and N.Z. conceptualised the study. All authors reviewed the manuscript.

## Acknowledgements

The authors acknowledge the support by the High Performance and Cloud Computing Group at the Zentrum für Datenverarbeitung of the University of Tübingen and the Federal Ministry of Education and Research (BMBF) through grant no. 031 A535A. We thank the Interfaculty Institute for Biomedical Informatics (IBMI) at the University of Tübingen for providing the computational resources essential for this study. NGS sequencing methods were performed with the support of the DFG-funded NGS Competence Center Tübingen (INST 37/1049-1). Additionally, we thank the Deutsche Forschungs-gemeinschaft (DFG, German Research Foundation) under Germany’s Excellence Strategy—EXC 2124—390838134 for structural support.

