## Supplementary Figures for "Ultra-deep long-read metagenomics captures diverse taxonomic and biosynthetic potential of soil microbes"

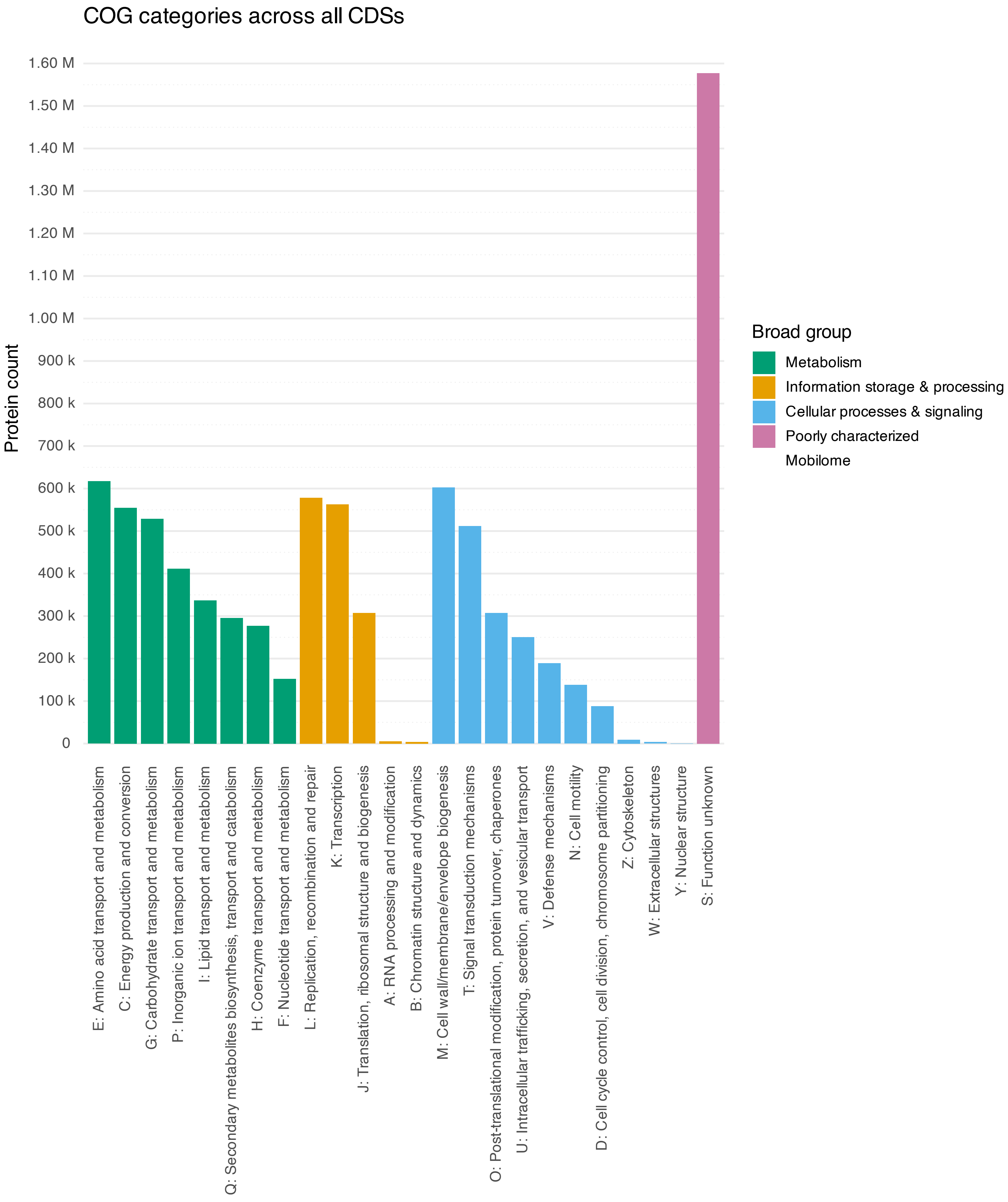


Supplementary Figure 1: Distribution of predicted proteins into COG categories.

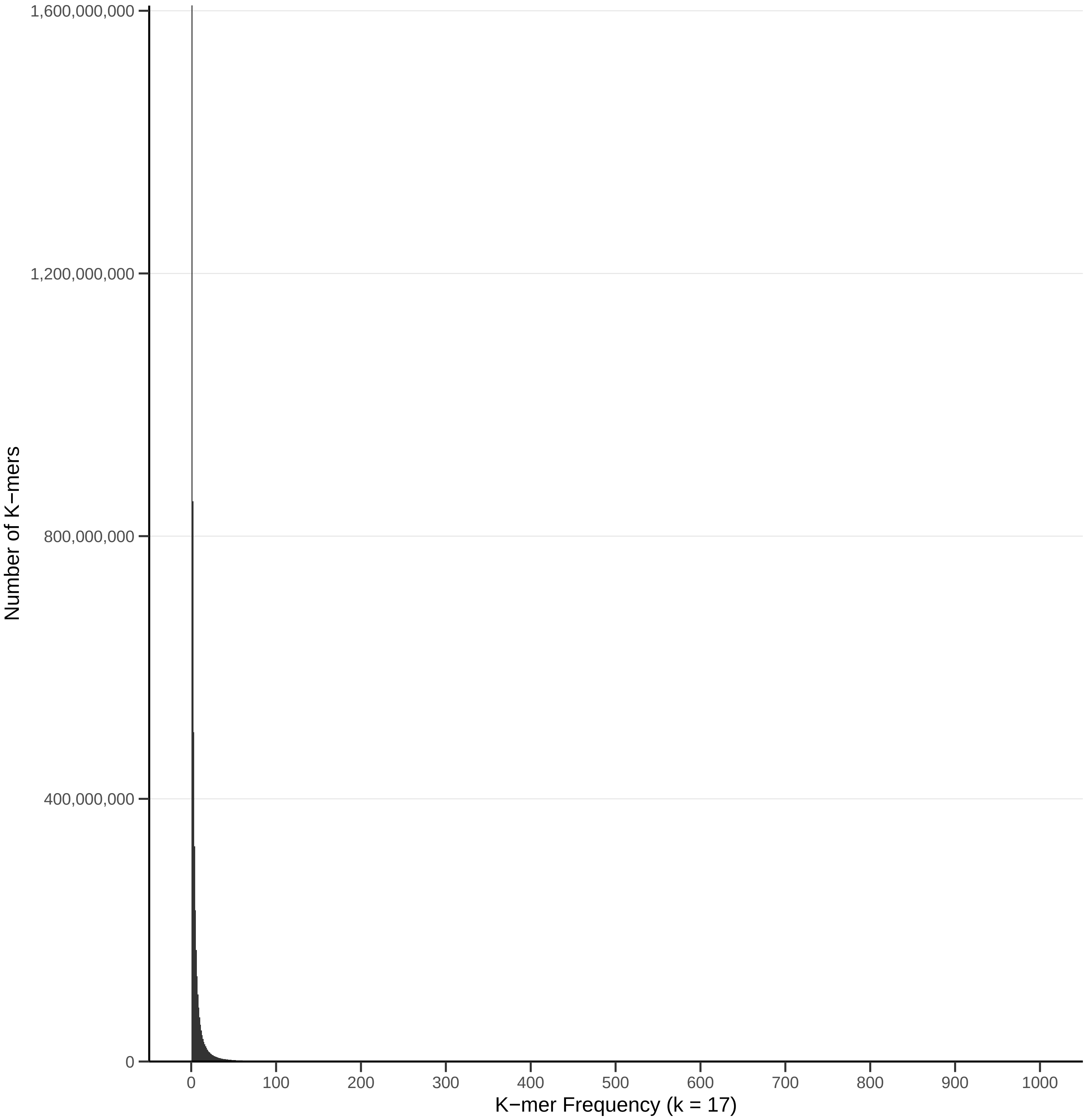


Supplementary Figure 2: 17-mer spectrum of Illumina reads.

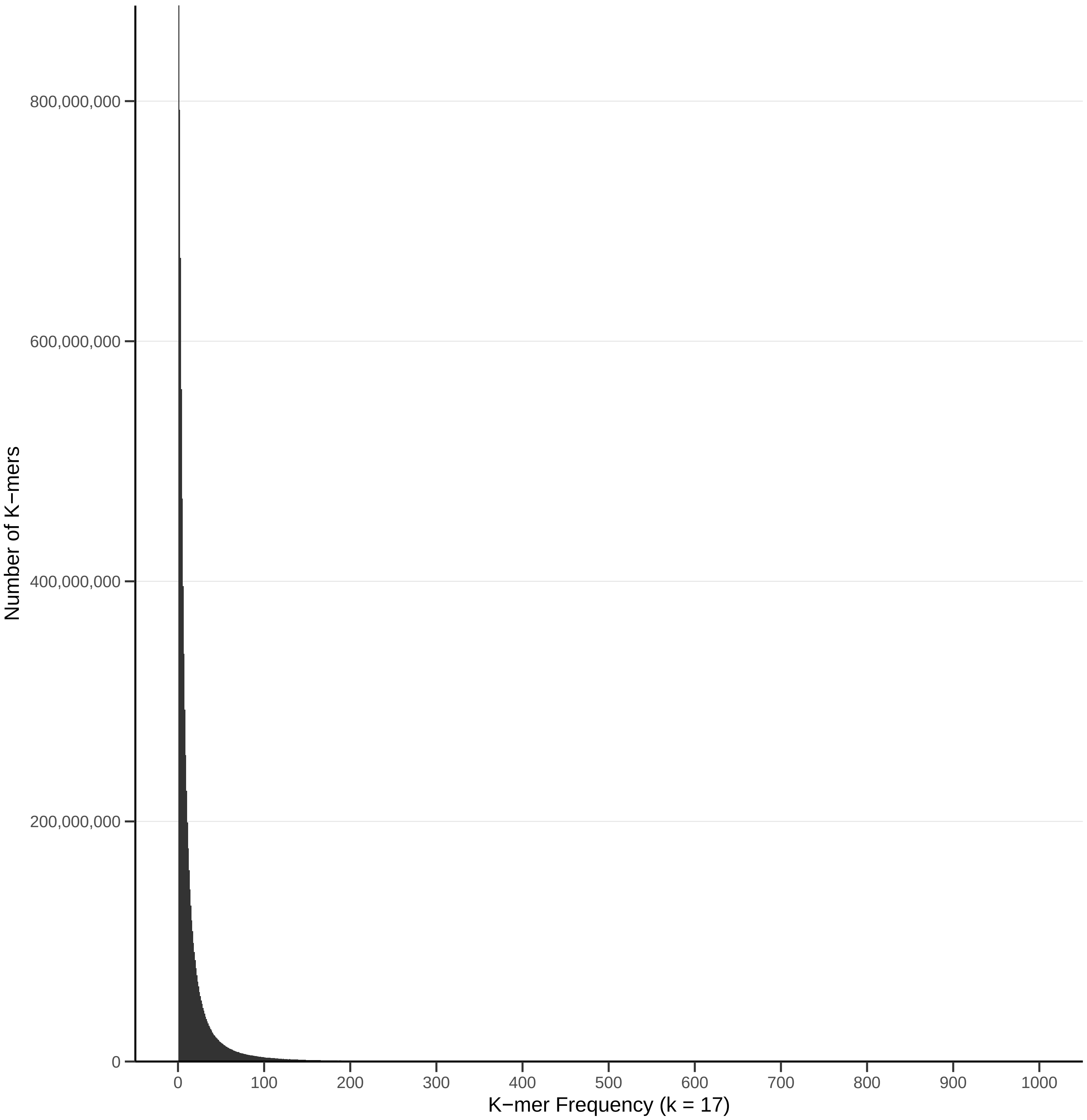


Supplementary Figure 3: 17-mer spectrum of Nanopore reads.

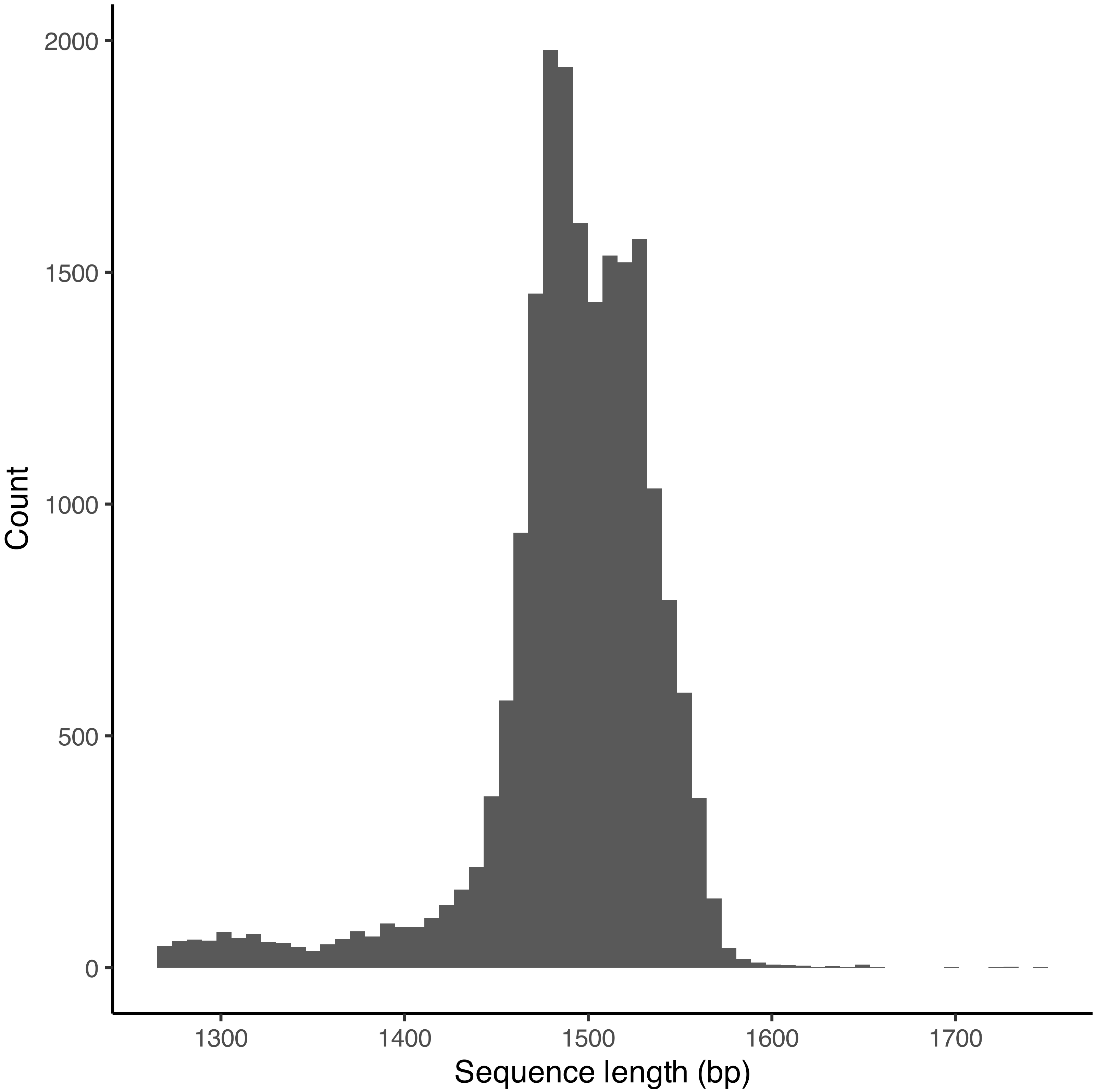


Supplementary Figure 4: Length distribution of the 16S rRNA gene sequences extracted from raw Nanopore reads.

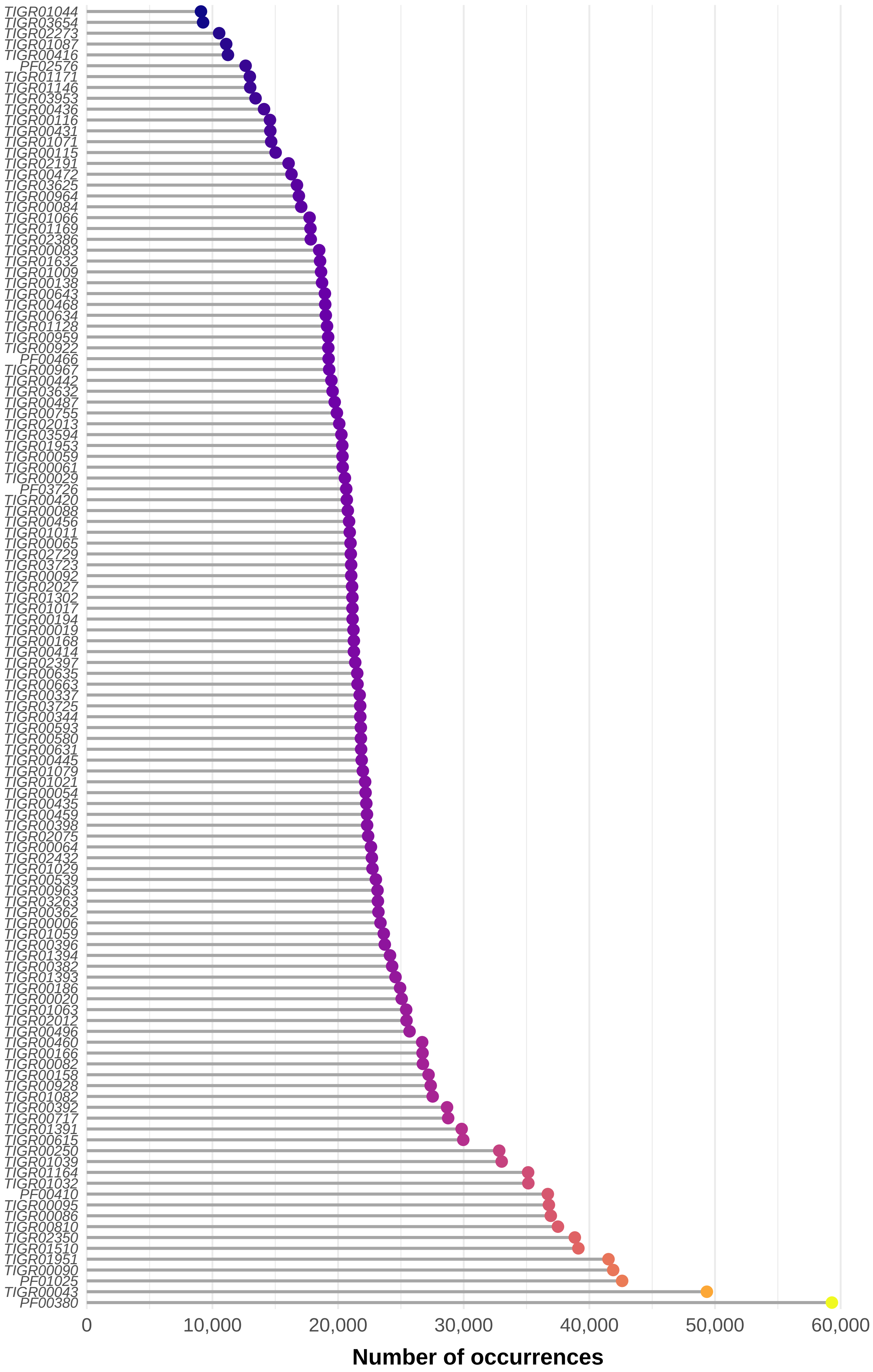


Supplementary Figure 5: The number of occurrences of each marker gene from the bac120 set in raw Nanopore reads.
